# Composition of the Lipid Droplets of the Bovine Corpus Luteum

**DOI:** 10.1101/2020.02.13.948117

**Authors:** Heather A. Talbott, Michele R. Plewes, Crystal Krause, Xiaoying Hou, Pan Zhang, William B. Rizzo, Jennifer R. Wood, Andrea S. Cupp, John S. Davis

## Abstract

Establishment and maintenance of pregnancy is dependent on progesterone synthesized by luteal tissue in the ovary. Our objective was to identify the characteristics of lipid droplets (LDs) in ovarian steroidogenic cells. We hypothesized that LDs are a major feature of steroidogenic luteal cells and store cholesteryl esters. Bovine luteal tissue was used for whole tissue analysis. Further analyses were performed on isolated ovarian steroidogenic cells: granulosa and theca cells of the follicle, and small/large luteal cells. Isolated luteal LDs were collected for lipid/protein analyses. Luteal tissue contained perilipins 2/3/5, hormone-sensitive lipase and abhydrolase domain containing 5. Luteal tissue was enriched in TGs compared to other tissues, except of adipose tissue. Large and small luteal cells were distinguished from follicular cells by the presence of LDs and LD-associated proteins. Furthermore, LDs from large luteal cells were numerous and small; whereas, LDs from small luteal cells were large and less numerous. Isolated LDs contained nearly all of the TGs and cholesteryl esters present in luteal tissue. Isolated luteal LDs were composed primarily of TG, with lesser amounts of cholesteryl esters, diglyceride and other phospholipids. Bovine luteal tissue LDs are distinct from LDs in other bovine tissues, including follicular steroidogenic cells.

## 3. Introduction

Luteal tissue forms in the ovary during each estrus or menstrual cycle and synthesizes progesterone, a steroid critical for early embryonic development and survival during pregnancy (1, 2). Luteal tissue has a tremendous ability to synthesize progesterone, secreting up to 40 mg/day in humans (3), and even greater quantities in cattle (4). The majority of the cholesterol utilized for progesterone biosynthesis comes from the blood in the form of high density lipoprotein-derived cholesteryl esters with smaller amounts from low density lipoprotein (5). Lipoproteins are internalized either through receptor-mediated endocytosis or selective cellular uptake, where cholesterol is sorted from lipoproteins within endosomes (5). Endosomal cholesterol is then believed to be trafficked to mitochondria for immediate progesterone biosynthesis or stored as cholesteryl esters in lipid reservoirs, also known as lipid droplets (LDs) (5, 6). In addition to its vital role in fertility in the mammal, progesterone is an essential precursor of androgens, estrogens, glucocorticoids and mineralocorticoids. Therefore, the high steroidogenic output of luteal tissue allows for detailed studies of mechanisms of steroidogenesis, which are likely conserved among all steroidogenic cells.

LDs are unique organelles that store neutral lipids and are coated with LD-associated proteins that are embedded within the surrounding phospholipid monolayer. These LD-associated proteins stabilize the LD, interact with additional proteins that insert or remove lipids from the LD core, enable LD trafficking, and mediate association of LDs with other organelles (7). The perilipin (PLIN) proteins, designated PLIN1-5, are a family of LD coat proteins that are important for stabilizing LD structure and provide a platform for protein assembly on the LD surface (7). Although LDs have been observed in nearly all tissues, LDs have been most extensively studied in adipose cells, where they form large unilocular droplets (7). In many circumstances, the formation of LDs is a sign of pathological conditions, such as the cholesteryl ester-laden foamy macrophage in atherosclerotic lesions (8), fatty liver disease due to liver damage (9, 10), or adiposity due to storage of excess lipids (11). However, in steroidogenic tissues such as the ovary, LDs are a prominent and generally non-pathogenic feature, except in cases of genetic disorders of steroidogenesis (12). The LDs in steroidogenic tissues are proposed to store cholesteryl esters that are used for steroidogenesis (6, 13).

Ovarian luteal tissue is easily distinguished from other tissues due to the abundant lipid content and cytoplasmic LDs. Luteal LDs exist in all species examined to date including: mice (14), rats (15), sheep (16), cattle (17), pigs (18), buffalo (19), rabbits (20), bats (21), non-human primates (22), and humans (23). The presence of LDs is used to distinguish active steroid-secreting luteal cells and are reported to contain both cholesterol and cholesteryl esters that can be used to synthesize steroids in both humans and cattle (24, 25). Strikingly, in rats and rabbits, luteal LDs are primarily composed of cholesteryl ester, and stimulation of steroid synthesis reduces LD cholesteryl ester concentrations (26). Luteal LDs are known to be regulated by diet (27), luteal trophic hormones like, luteinizing hormone, which decreases LDs (26) and luteal lytic hormones, like prostaglandin F2α, that increase LDs (28). With the knowledge of LD-associated proteins and lipids in other tissues, we set out to characterize the LD features in the bovine luteal tissue, while it was actively secreting progesterone. The present study compares luteal LDs to LDs in other tissues, and provides the first characterization of LDs in large and small steroidogenic luteal cells, and provides a comprehensive analysis of the phospholipid and neutral lipid composition of luteal LDs.

## 4. Materials and Methods

### 4.1. Animals

Female cattle (n = 15) of composite breeding were synchronized using two intramuscular injections of prostaglandin F2α (25 mg; Lutalyse®, Zoetis Inc., Kalamazoo, MI) 11 days apart. At day 3 or day 10 after ovulation, 3-5 cows were subjected to a bilateral ovariectomy through a right flank approach under local anesthesia, as previously detailed (29, 30). Luteal tissue was prepared for microscopy and remaining tissue was frozen using liquid N2. Day 10 corpora lutea weighed significantly more than day 3 corpora lutea (4.7 ± 0.46 vs. 2.8 ± 0.65 g, respectively). However, in all other measures examined in the course of this study there were no significant differences. Therefore, data from both day 3 and day 10 were pooled to provide additional statistical power and narrow the confidence intervals associated with progesterone secreting bovine corpora lutea. All animal procedures were completed under an IACUC-approved protocol and performed at the University of Nebraska—Lincoln, Animal Sciences Department.

### 4.2. LD staining in luteal tissue

Tissue sections were frozen in OCT (Tissue-Tek) and transported on dry ice. Frozen samples were kept at −80 °C until sectioning using a Leica CM3050S instrument and attached to silane-coated slides before fixation in 10% phosphate-buffered formalin for 1 h. Select fixed slides were stained with oil red O and counterstained with Harris’ hemotoxin using an automated slide staining set up at the University of Nebraska Medical Center Tissue Sciences Facility. Slides were scanned at 40x using Ventana’s Coreo Au Slide Scanner. Images were analyzed by Definiens Tissue Studio (Munich, Germany) to quantify nuclei number and area occupied by oil red O.

Coronal sections (through the stomata) of luteal tissue were fixed in 3% (w/v) paraformaldehyde and 0.2% glutaraldehyde in PBS, pH 7.4, post-fixed in 2 % OsO4, resin-embedded, and ultra-thin sectioned for electron microscopy. Transmission electron microscopy (TEM) images were captured using a Hitachi H7500 at the University of Nebraska Lincoln Center for Biotechnology. Three images (magnification: 8,000x) from luteal tissue from each animal were used for quantification of LD number and area using ImageJ (31).

Additional fixed luteal tissue slides were immunolabeled for a marker of steroidogenic cells, Hydroxy-Delta-5-Steroid Dehydrogenase, 3 Beta- And Steroid Delta-Isomerase 1 (HSD3B), and incubated at 4 °C for 24 h. Slides were washed 3 times with 0.1% Tween in PBS to remove unbound antibody and counterstained using appropriate secondary antibodies, BODIPY 493/503 and 4′,6-diamidino-2-phenylindole (DAPI). Slides were washed again and mounted to glass microscope slides using Fluoromount-G (Electron Microscopy Sciences) and stored at −22 °C until imaging. Additional antibody and staining information are available in Error! Reference source not found..

### 4.3. Isolation of granulosa, theca, large luteal, and small luteal cells

For luteal cell preparations, bovine ovaries were collected during early pregnancy, dissociated with collagenase, and elutriation was performed to prepare enriched preparations of small and large luteal cells, as described previously (32, 33). The average purity of small luteal cells was > 90% and large luteal cells was > 50%.

Follicular granulosa and theca cells were prepared from bovine ovaries with large follicles (> 8 mm diameter). Follicular fluid was removed with a 20-gauge needle, the needle was re-inserted, and the granulosa cells were suspended in an equal volume of DMEM-F12 culture media. After the granulosa cells were removed, the theca interna was removed with fine forceps. Granulosa cells were washed by centrifugation three times at 150 rcf for 5-10 min and filtration through a 70 μm nylon mesh. The theca interna were suspended in 103 IU/mL collagenase 2 (Atlanta Biologicals) in DMEM/F12 and dispersed using constant agitation at 37 °C for 1 h. Dispersed theca cells were removed from the undigested tissue by filtration through a 70 μm mesh then washed by centrifugation three times at 150 rcf for 5-10 min.

### 4.4. Microarray

Bovine gene expression arrays from NCBI GEO repository (GSE83524) were minded to analyze expression of expression of LD components in freshly isolated bovine granulosa (GC, n = 4) and theca (TC, n = 3) cells from large follicles and from purified preparations of bovine small (n = 3) and large (n = 3) luteal cells from mature corpora lutea. Details of the isolation and analysis were previously published (34).

### 4.5. LD Isolation from Tissue

Bovine luteal, adipose, cardiac, hepatic and pulmonary tissues were obtained from a local abattoir and transported to the laboratory in cold BPS. The tissues (∼2.5 g) were washed thoroughly in TE buffer (10 mM Tris, 1 mM EDTA, pH 7.4). Tissue was minced in 10 mL tissue homogenate buffer (60% sucrose w/v in TE buffer containing protease and phosphatase inhibitor cocktail) and homogenized with a Teflon Dounce homogenizer in a glass vessel. The post-nuclear supernatant fraction was obtained after centrifugation at 2000 rcf for 10 min. The supernatant was loaded into a 30 mL ultracentrifuge tube and overlaid sequentially with 40%, 25%, 10%, and 0% sucrose w/v in TE buffer containing protease and phosphatase inhibitor cocktails. Samples were centrifuged at 110,000 rcf for 30 min at 4 °C with no brake in a Beckman Coulter Avanti J-20 XP ultracentrifuge using an SW 32 Ti rotor. LDs concentrated in a band at the top of the gradient were harvested and concentrated by centrifugation at 2000 rcf for 10 min at 4 °C. The protocol was modified from (35, 36).

### 4.6. Western Blots

Western blots were performed as previously described (32) with the antibodies and reagents described in Error! Reference source not found..

### 4.7. Lipidomics

Lipids from luteal LDs were extracted using a standard Bligh and Dyer extraction protocol (37) and then dried under a stream of nitrogen and sent to Avanti Polar Lipids for lipidomic profiling of free sterols, cholesteryl ester, TG, DGs, phospholipids, and sphingolipids. The molecular species within each class were identified, quantified and summed to report the average lipid profile of bovine luteal LDs. To provide resolution and quantitative ability beyond the mass resolution of the tandem quadrupole mass spectrometers employed, molecular species were resolved by reversed-phase liquid chromatography in the presence of class and sub-class specific internal standard compounds added to each sample. The compounds were detected by tandem MRM MS/MS for mass specific fragment ions according to lipid class and molecular weight of the compound. Selectivity was further enhanced by scheduling the detection of each compound according to its elution from the HPLC column, known as scheduled MRM. The semi-quantization was calculated using the integrated area of each analyte’s MRM peak, divided by the appropriate internal standard peak area, and multiplied by its known concentration. Quantification of cholesterol and cholesteryl esters were directly calculated with standards and internal standards from calibration response curves. Lipid concentrations were normalized to the corresponding protein concentration of each sample and as a mol % relative to lipid class.

### 4.8. High Performance Thin Layer Chromatography

For lipid analyses, cell suspensions were extracted with chloroform-methanol (1:1). HPTLC was performed as previously described with modification to a single solvent system [petroleum ether (b.p. 60-70 °C)-ethyl ether-acetic acid (45:5:0.5] (38). The images were analyzed using UVP Vision Works LS software by calculating the area under the curve after lane-specific straight line background correction. A mixture of the following standard lipids was co-chromatographed: cholesterol, trioleate glyceride, cholesterol palmitate, and oleic acid. Preliminary analyses were completed to establish the linearity of detection for each lipid class to ensure that lipids did not exceed the linear range for quantitation. For every plate of cellular lipids, five lanes of varying amounts of lipid standards were simultaneously run to generate standard curves for quantitation. The amount of each cellular lipid was expressed as μg lipid/mg cell protein or μg lipid/mg initial tissue mass. The protocol was adapted from (39).

### 4.9. Confocal microscopy and analysis

To characterize LDs in small and large bovine luteal cells, thin-layer cell preparations for confocal microscopy of enriched small luteal cells and large luteal cells was performed using Cytofuge 2 (Beckman Counter). Cells were then fixed with 10% neutral buffered formalin at 4 °C for 30 min. Cells were stained with BODIPY 493/503 and phalloidin (Error! Reference source not found.) for 1 h at room temperature and mounted using Fluoromount-G.

To determine the colocalization of BODIPY C12 and TopFluor Cholesterol in luteal LDs, **e**nriched small (5×10^4^ cells/cm^2^) or large (2×10^4^ cells/cm^2^) luteal cell cultures were seeded onto **s**terile No. 1 glass coverslips (22 × 22 mm) and cultured as previously described. Cells were treated with TopFluor Cholesterol and BODIPY C12 for 48 h to allow incorporation in LDs. Cells were then incubated in Mitotracker Deep Red for 45 min to label mitochondria (Error! Reference source not found.). Cells were then fixed with 4% paraformaldehyde and at 4 °C for 30 min and mounted using Fluoromount-G. Images were collected using a Zeiss confocal microscope equipped with a 60× oil immersion objective (1.4 N.A) and acquisition image size of 512 × 512 pixel (33.3 μm × 33.3 μm). Cells were randomly selected from each slide and 0.33 μm slice z-stacked images were generated from bottom to top of each cell. A 3-dimensional image of each cell was created, and area of individual cell was generated using Zen software. Images were converted to maximum intensity projections and processed utilizing ImageJ (National Institutes of Health) analysis software. LD size and number were quantified with ImageJ using AnalyzeParticles function in threshold images, with size (square pixel) setting from 0.1 to 100 and circularity from 0 to 1. Outputs were then converted into microns. For colocalization of BODIPY C12 and TopFluor Cholesterol, z-stack images were analyzed in Image J using the JACoP plugin as previously described (40).

### 4.10. Statistical Analysis

All data are presented as the means ± SEM. Data was evaluated for normal distribution and log transformed if necessary. Specifics of statistical testing are described in the relevant figure legends. Statistical analysis was performed using GraphPad Prism (GraphPad Software, Inc) except for the microarray data which was analyzed as previously described (34).

## 5. Results

### 5.1. Visualization and quantification of luteal tissue LDs

Functional bovine luteal tissue prominently featured LDs, which occupied the majority of luteal tissue consistent with steroidogenic cell distribution within luteal tissue. However, vasculature and connective tissue invaginations within the luteal tissue (composed of non-steroidogenic endothelial and fibroblast cells) contained few LDs (**Figure 1A**). Luteal LDs, stained with oil red O, occupied an average area of 29.9 ± 8.56 μm^2^ per nucleus within luteal tissue sections (**Figure 1A & B**). Luteal LDs were an abundant ultrastructural feature of functional luteal tissue and were often in close proximity with mitochondria (**Figure 1 C)**. Individual LDs occupied an average area of 0.41 ± 0.04 μm^2^ in luteal tissue corresponding to an average diameter of ∼0.72 μm and ranged from 0.16-1.8 μm^2^ (**Figure 1 C & D**). Moreover, BODIPY-labeled LDs co-localized with cells immuno-labeled with a marker of steroidogenic cells, HSD3B, which confirmed luteal LDs are present within the steroidogenic cells of luteal tissue (**Figure 1E**).

**Figure 1.**
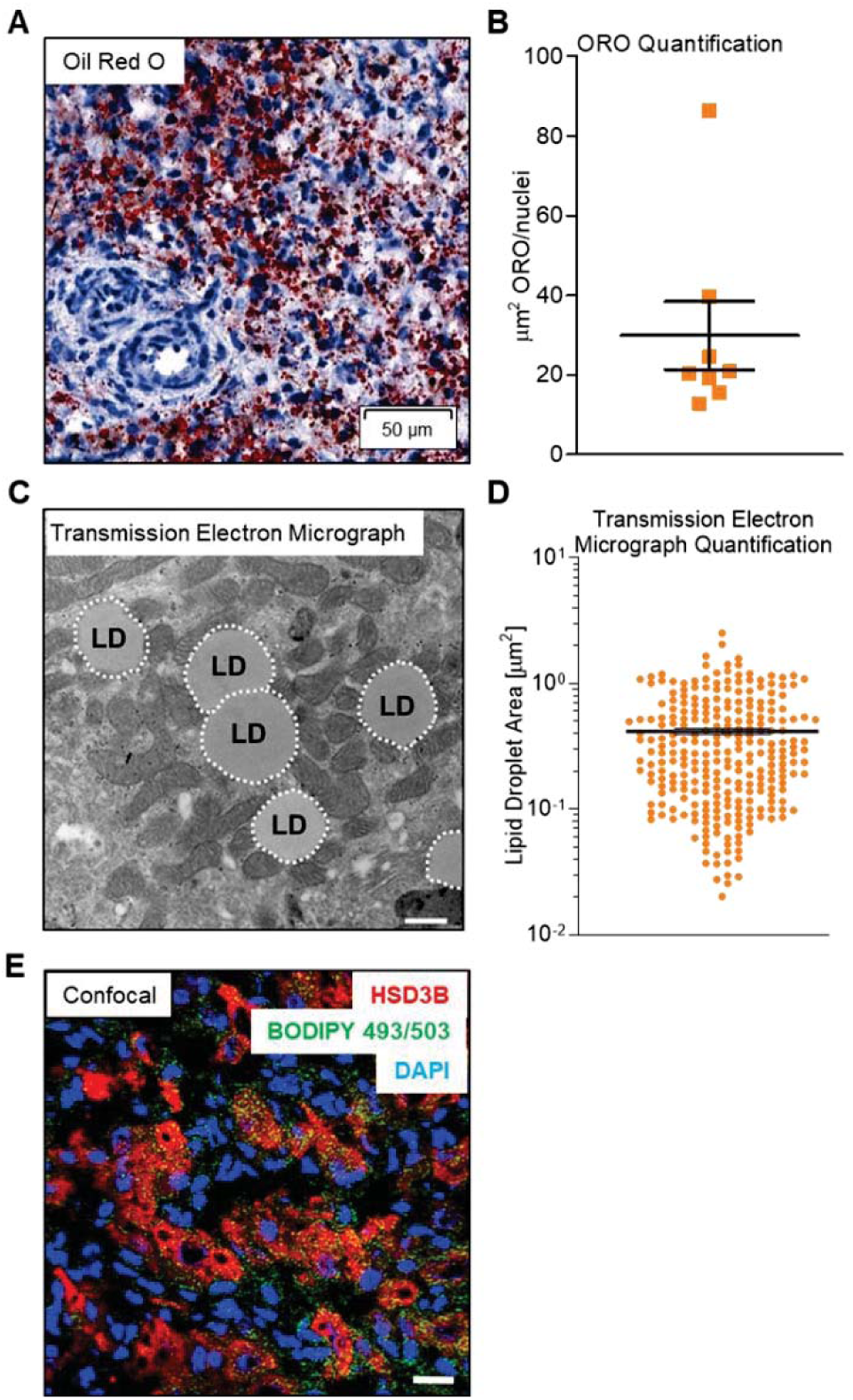
Visualization of luteal LDs. Luteal LDs were visualized using oil red O staining, transmission electron micrographs, and confocal microscopy. **Panel A**: Oil red O staining (red) of lipids in frozen tissue sections of functional bovine corpus luteum counterstained with hematoxylin (blue). Representative image shown at 40x, scale bar = 50 μm. **Panel B**: Automated quantification of tissue area occupied by oil red O staining, each point demonstrates the area (μm^2^) occupied by red immunohistochemistry staining in three randomly chosen images per animal (n = 6, mean ± SEM indicated with black lines). **Panel C**: Transmission electron micrograph of bovine luteal tissue demonstrating multiple LDs (labeled LD) surrounded by mitochondria. Representative image shown at 21,000x, scale bar = 500 nm. **Panel D**: Logarithmic graph of the quantification of area (μm^2^) occupied by individual LDs in three randomly chosen images per animal (n = 6, mean ± SEM are indicated with a black line). **Panel E**: Representative confocal image obtained from frozen tissue sections of functional luteal tissue co-labeled with BODIPY493/503 (green; LD), HSD3B (red; marker for steroidogenic cells), and 4′,6-diamidino-2-phenylindole (DAPI, blue). Representative image shown at 63x, scale bar = 20 μm.

### 5.2. Luteal tissue compared to other LD-associated tissues

Luteal tissue had a distinct composition of LD-associated proteins compared to other tissues (**Figure 2A**). Luteal tissue had considerable PLIN2 and PLIN3 protein, little to no PLIN1, and a moderate amount of PLIN5. In contrast, visceral adipose tissue had abundant PLIN1, hepatic tissue expressed both PLIN2 and PLIN3, and cardiac tissue had high amounts of PLIN5. Bovine luteal tissue also had substantial amounts of lipid-modifying enzymes hormone sensitive lipase and abhydrolase domain containing 5(ABHD5) and lesser amounts of adipose triglyceride lipase (ATGL) and sterol O-acyltransferase (SOAT1). Hepatic tissue had the most SOAT1 protein and also expressed ABHD5, but had little ATGL or hormone sensitive lipase. Adipose tissue had the most hormone sensitive lipase, large amounts of ATGL, and less SOAT1 than cardiac or hepatic tissue. Finally, cardiac tissue had large amounts of ATGL, intermediate amounts of SOAT1, and a small amount of hormone sensitive lipase. Luteal tissue additionally had a unique lipid composition, which is particularly evident in the high TG content compared to pulmonary, hepatic, and cardiac tissue (**Figure** 2**B**). Adipose tissue, as expected, had nearly 100-fold more TG than other tissues. Cholesteryl esters were a minor lipid class in all tissues investigated, and cholesteryl ester levels were nearly absent in bovine cardiac tissue. Sterol abundance was lowest in cardiac tissue and highest in adipose tissue; whereas, free fatty acids were highest in adipose tissue (4-fold), followed by hepatic tissue.

**Figure 2.**
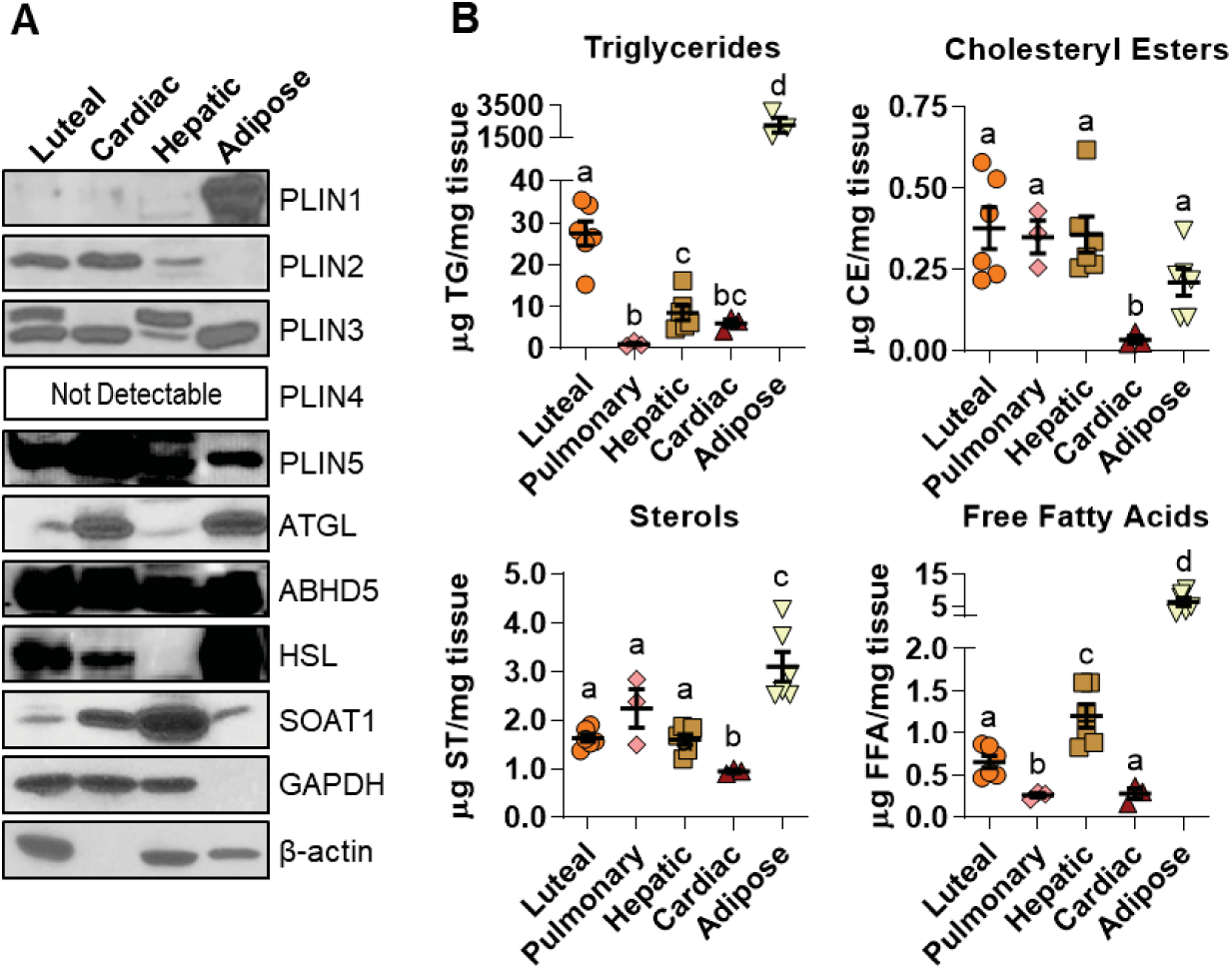
Lipid droplet (LD)-associated protein and lipid content of luteal tissue. Bovine luteal, cardiac, hepatic, pulmonary and visceral adipose tissues were collected to compare LD-associated protein and lipid content of luteal tissue between LD associated tissues. **Panel A**: The presence of the LD-coat proteins (PLIN1-5), key neutral lipid hydrolysis enzymes (ATGL, ABHD5, HSL), lipid forming enzymes (SOAT1) and loading controls (β-actin & GAPDH) were assessed by western blot. **Panel B**: Lipid content of luteal tissue (n = 6) in comparison to pulmonary (n = 3), hepatic (n = 6), cardiac (n = 3), and visceral adipose tissue (n = 6) was determined by HPTLC analysis to assess relative amounts of major neutral lipid classes. Horizontal line and error bars indicate mean ± S.E.M, significance was determined using mixed-effects model with matching lipid measurement within each sample after log transformation of values and Tukey’s multiple comparisons test. Means with different letters differ significantly between tissues (*P* < 0.05).

### 5.3. Comparison of follicular and luteal cell type LD features

The follicular granulosa and theca cells differentiate to give rise to large and small luteal cells after ovulation (41). LD content was compared between the follicular and luteal cell types. Granulosa and theca cells had fewer and smaller LDs than the luteal cell types, as assessed by BODIPY staining of LDs and confocal imaging (**Figure 3A**). Cytoplasmic staining for aromatase, CYP19A1, confirmed the identity of granulosa cells. A microarray comparison (34) between granulosa, theca, large luteal cells and small luteal cells indicated that mRNA abundance of LD-associated proteins *PLIN2, PLIN3* and hormone sensitive lipase (*LIPE)* were increased in luteal cell types compared to follicular cells (**Figure 3B**). Similarly, luteal tissue had greater amounts of hormone sensitive lipase and PLIN2 protein compared to granulosa and theca cell isolates. (**Figure** 3**C**). The lipid content of isolated granulosa, theca, large luteal cells, and small luteal cells were compared using HPTLC. Large and small luteal cells had more TG than granulosa or theca cells (**Figure 3D**). Granulosa cells contained more TG, sterols, and free fatty acids than theca cells. Surprisingly, cholesteryl ester concentration tended to be highest in granulosa cells.

**Figure 3.**
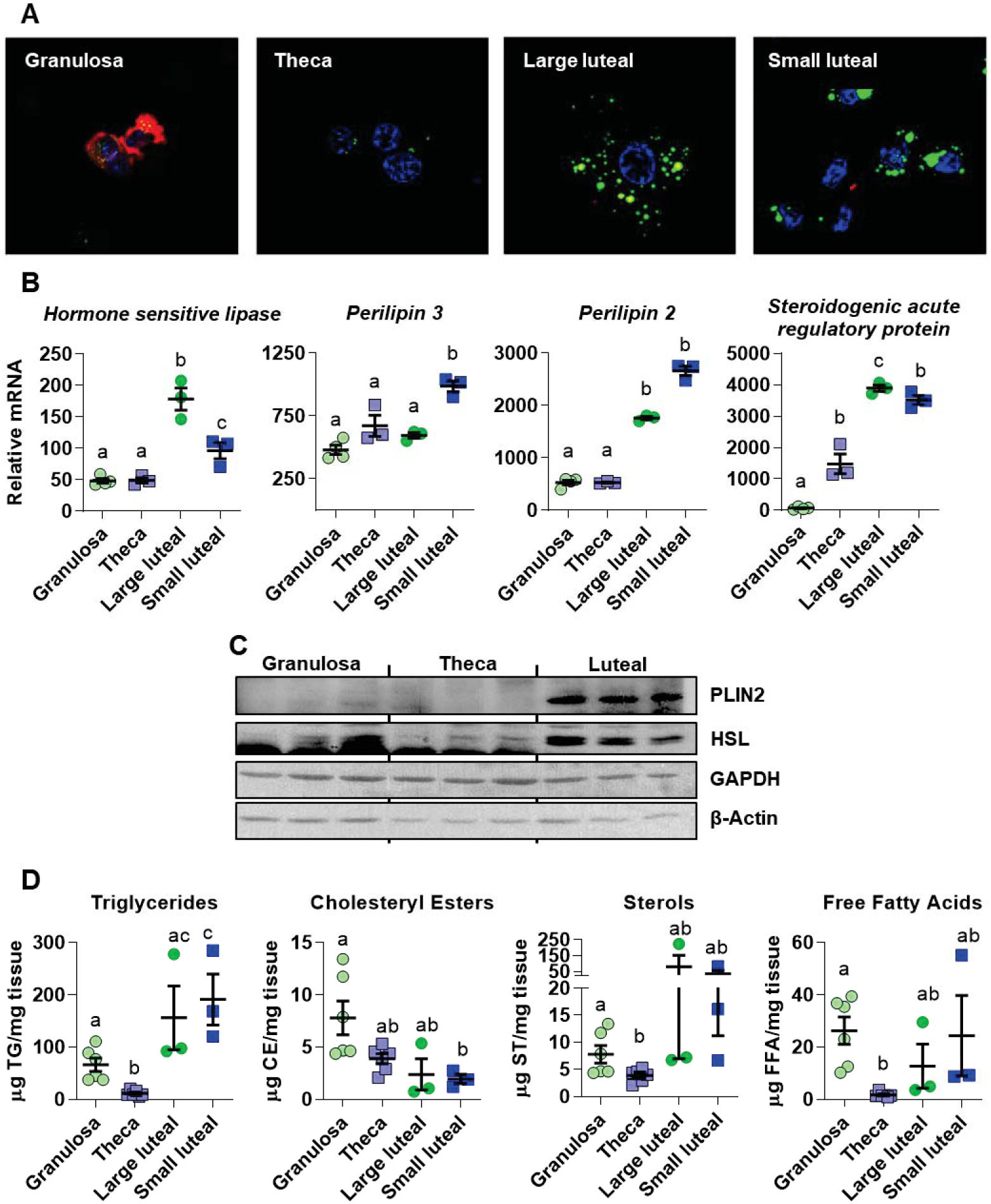
Comparison of follicular and luteal LD properties. Follicular granulosa cells, theca cells, large luteal cells, and small luteal cells were isolated from bovine ovaries as described in the Methods. **Panel A**: Confocal fluorescent image showing LD staining in freshly isolated bovine granulosa, theca, large luteal, and small luteal cells. LDs were stained with the neutral lipid dye BODIPY 493/503 (green), and cells were immuno-labeled using an aromatase antibody to specifically label granulosa cells (red), and the nuclei are counter-stained with 4′,6-diamidino-2-phenylindole (DAPI, blue). All images are equal magnification. **Panel B**: Microarray analysis of mRNA abundance of LD-coat proteins: perilipin 2 and 3, cholesteryl esterase: hormone sensitive lipase, and steroidogenic marker: steroidogenic acute regulatory protein. Significance determined in (34). Means with different letters differ significantly between cell types (*P* < 0.05). **Panel C**: Western blot of LD-associated proteins in follicular granulosa and theca cells in comparison to luteal tissue. **Panel D**: HPTLC analysis of freshly isolated granulosa (n = 6), theca (n = 6), large luteal (n = 3) and small luteal cells (n = 3). Means ± SEM overlay individual measurements, significance was determined using two-way ANOVA and Tukey’s multiple comparisons test with matching lipid measurements within each sample after log transformation of values. Means with different letters differ significantly between cell types (*P* < 0.05).

Other groups (42) have suggested cholesterol and TG can segregate into distinct LD populations, therefore large and small luteal cells were incubated with Topfluor Cholesterol and the fatty acid BODIPY C12 and imaged by confocal microscopy. TopFluor Cholesterol and BODIPY C12 colocalized in both large luteal cell (92.7 ± 1.2%) and in small luteal cells (91.3 ± 1.2%) (**Figure 4AB**). Isolated large and small luteal cells were imaged using confocal microscopy to evaluate LD properties. The large luteal cells had significantly more LDs (271 ± 22) compared to small luteal cells (89 ± 8) (Figure 4C **& E**); whereas, small luteal cell LDs were significantly greater in volume (695.5 ± 112.3 nm^3^) than large luteal cell LDs (293.2 ± 39 nm^3^) (Figure **B, C & E**); these measurements corresponded to an average LD diameter of 1.1 μM for small luteal cells and 0.82 μM for large luteal cells. Small luteal cells had more lipid content than large luteal cells measured using mean fluorescence intensity to estimate the total lipid content (**Figure 4D**).

**Figure 4.**
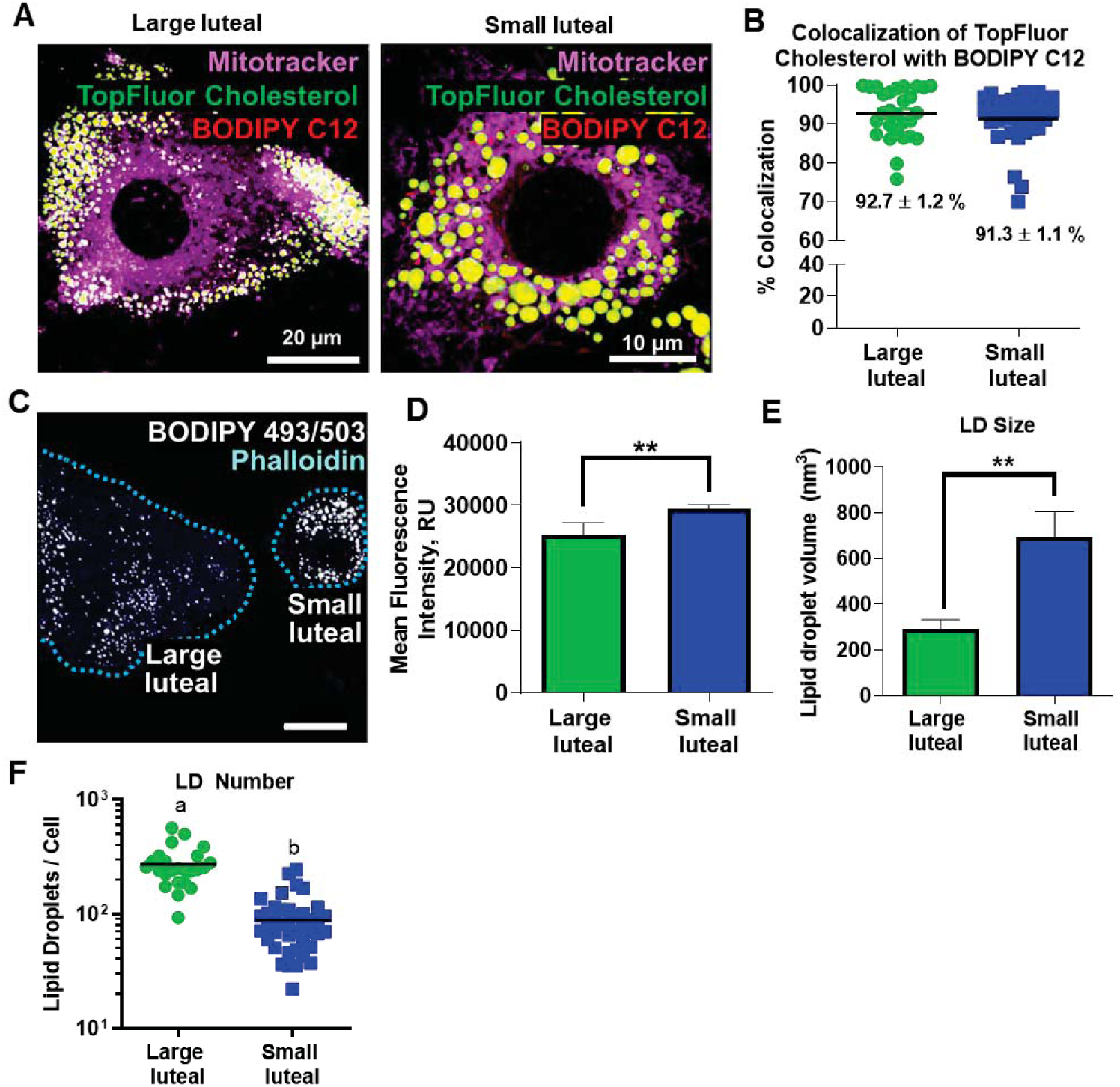
Differences in LDs between small and large luteal cells. Individual LD measurements in freshly isolated large luteal cells and small luteal cells were determined using confocal microscopy. **Panel A:** Representative confocal micrographs of enriched large luteal and small luteal populations pre-loaded with BODIPY C12 and TopFluor Cholesterol for 48-h. Mitochondria (Mitotracker; magenta), TopFluor Cholesterol (green), and BODIPY C12 (red); Micron bar represents 20 and 10 μm, respectively. **Panel B:** Quantitative analysis of percent colocalization of TopFluor Cholesterol with BODIPY C12 in large luteal cells (green) and small luteal cell (blue). **Panel C:** Individual lipid droplet (LD) measurements in freshly isolated large and small luteal cells were determined after BODIPY staining of three-dimensional rendering of confocal images. Representative images of large luteal cells and small luteal cells. BODIPY 493/503 (white) and Phalloidin (blue); Dashed blue line indicates cell boundaries. Micron bar represents 20 μm. **Panel D:** Quantitative analysis of mean fluorescence intensity for large luteal and small luteal cells. Significance of data was assessed with unpaired two-tailed t-test. **Panel E:** Quantitative analysis of individual LD size, large luteal (25 individual cells), small luteal cells (41 individual cells) from 3 animals are displayed as mean ± SEM. **Panel F**: Logarithmic graph displaying the quantitative analysis of number of LDs per cell, significance was assessed using unpaired two-tailed t-test after log transformation of the data.

### 5.4. Isolation and composition of luteal LDs

A step-wise sucrose gradient isolated LDs from total luteal tissue. Isolated luteal LDs were yellow in color in comparison to LDs from other tissues (**Figure 5A**). Imaging of isolated LDs by embedding and visualization through TEM demonstrated intact LDs are relatively free from other cellular organelles and debris when isolated using this protocol (**Figure 5B**). The lipid class composition of the total luteal tissue fraction, the isolated LD fraction, and the LD depleted fraction were compared using HPTLC to determine the cellular localization of the various lipid classes (**Figure 5C**). TGs were primarily stored in luteal LDs, with nearly no TG found in LD-depleted lysates. Similarly, cholesteryl esters were present in the LD fraction and absent in LD-depleted lysates. Sterols and free fatty acids showed little segregation between LDs and LD-depleted cellular fractions. Lipid extracts of LDs were characteristically yellow which deepened in color during concentration of LDs and of LD lipids. During HPTLC a yellow band, presumably carotenes, consistently migrated along the solvent front with cholesteryl esters (not shown). Additionally, a band of unknown lipid species consistently migrated between the TG and cholesteryl ester standards when separated by HPTLC (not shown).

**Figure 5.**
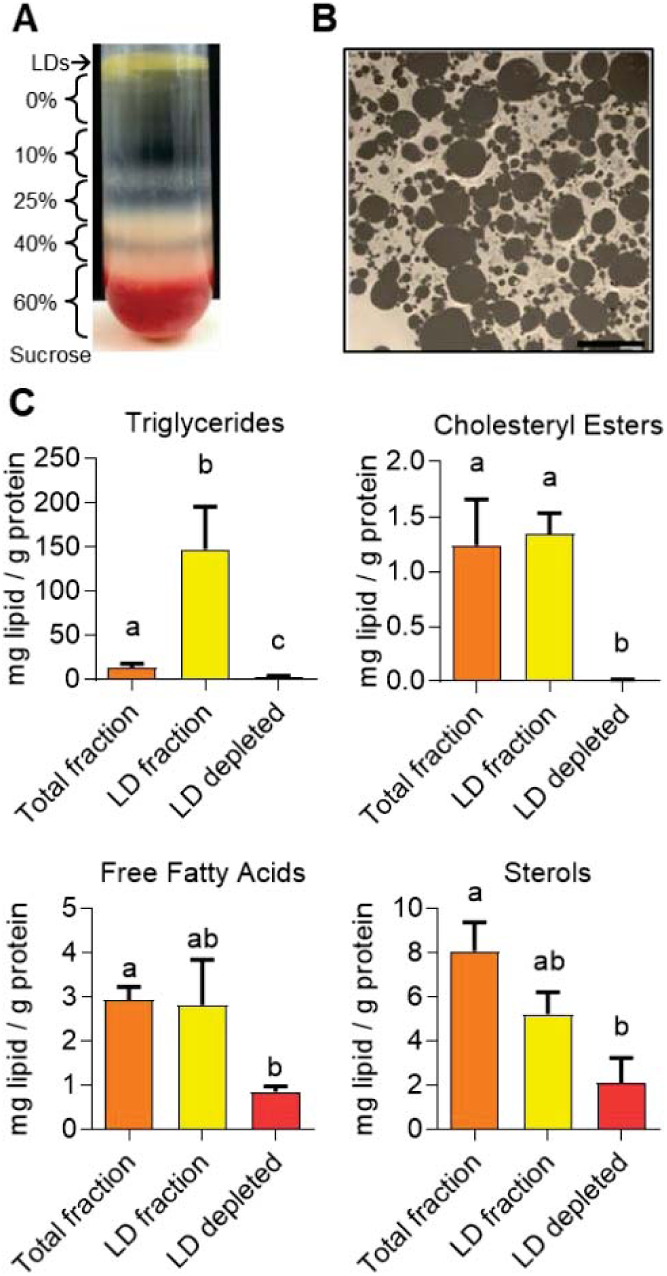
Isolated luteal LD properties. Sucrose gradient and ultra-centrifugation was used to isolate luteal LDs. Transmission electron micrography (TEM) images were obtained to validate successful isolation of luteal LDs. **Panel A**: Representative image of luteal LDs and sucrose gradient fractions following ultra-centrifugation. **Panel B**: Representative transmission electron micrograph from isolated LD preparations. Micron bar represents 6 μm. HPTLC was used to determine the lipid composition of luteal lipid droplets (LDs). Lipid analysis by HPTLC of whole luteal tissue fraction (total), LD fraction, and LD depleted post-nuclear supernatant. Lipid content of each fraction was normalized to protein content. Bars represent means ± SEM, significance was determined using mixed-effects model and Tukey’s multiple comparison test with matching lipid measurement within each sample after log transformation of values, n = 5. Means with different letters differ significantly between tissues (*P* < 0.05).

Lipidomics analysis of three different preparations of luteal tissue LDs confirmed that luteal LDs are primarily composed of TG (168 ± 42 pmol/μg protein, 89% of lipid class to total lipids) but also contain many other lipid classes (**Figure 6A**). Other neutral lipids included DG (5.7 ± 2.1 pmol/μg protein, 2.9%; and cholesteryl ester (3.65 ± 0.7 pmol/μg protein, 2.0%). Polar lipids were primarily composed of PC (3.9 ± .8 pmol/μg protein, 2.2%), SM (2.7 ± 0.3 pmol/μg protein, 1.5%), PI (1.7 ± 0.6 pmol/μg protein, 0.9%), PE (1.4 ± 0.4 pmol/μg protein, 0.8%) and PS (0.65 ± 0.04 pmol/μg protein, 0.4%). A number of other minor lipids representing less than 0.33 mol% of the total lipid pool were also detected including PG, lysophospholipids (LPI, LPC, LPE, LPS), Cer, GlucCer, and sphingoid bases (**Figure 6A**). Sterols were below the limit of detection (lowest concentration of standard curve) in all three LD samples. The polar to nonpolar lipid ratio was 7.12 ± 0.017, which corresponds to an average LD diameter of 282 nm if the properties of sphericity and surface area to volume ratios are used to calculate the expected LD diameter (35).

**Figure 6.**
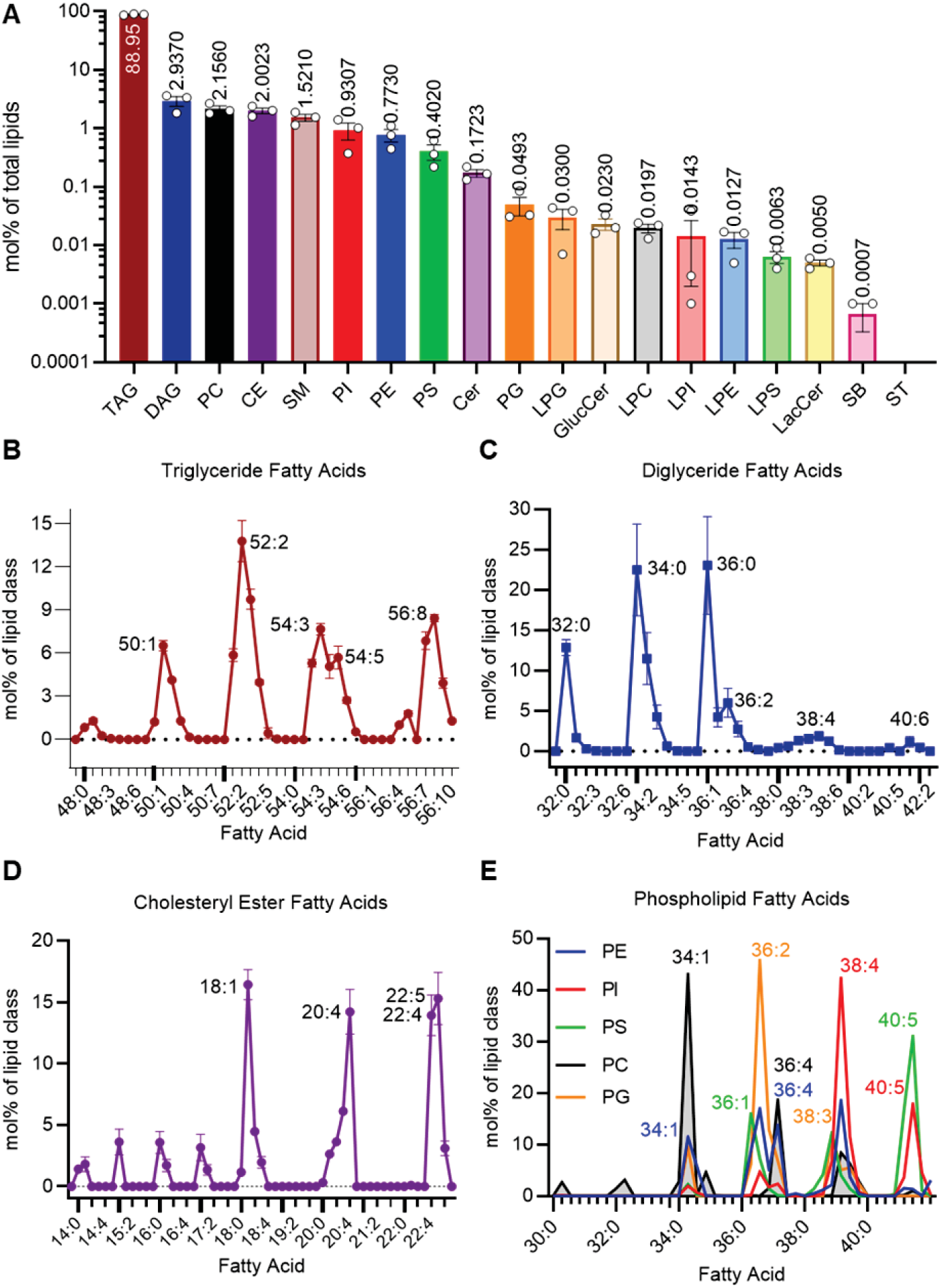
Lipidomic analysis of isolated luteal lipid droplets (LDs). LC/MS/MS with sMRM was used to detect lipid species with lipid class specific internal standards to allow for relative quantification. **Panel A:** The top 18 lipid classes present in bovine luteal LDs are shown from the largest proportion of the total lipid fraction to the smallest as measured as the mol% of the specific lipid class to the total lipid content, means ± SEM overlay individual measurements, n = 3. Cholesteryl Ester (CE); Glucosylceramide (GlucCer); sphingoid bases (SB). **Panel B:** The combined fatty acid composition of TG in luteal LDs are represented as the mol% within the TG class, with increasing degrees of saturation listed within the shortest to longest combined acyl chain length, peaks are labeled with the relevant fatty acid, points indicate means ± SEM. **Panel C:** The fatty acid composition of the diglycerides in luteal LDs are represented as the mol% within the diglyceride class, points indicate means ± SEM. **Panel D:** The combined fatty acid composition of the cholesteryl esters in luteal LDs are represented as the mol% within the cholesteryl ester class, points indicate means ± SEM. **Panel E:** The fatty acid composition of several phospholipid classes (PE PI, PS, PC and PG) in luteal LDs are represented as the mol% within the appropriate lipid class, peaks are labeled with the relevant fatty acid, inflection points indicate means.

Each lipid class was composed of a diversity of fatty acyl species. The TG class was represented by 28 species, of which 26 molecular species were found in all three samples (**Figure 6B**). Four principal groups with 52, 56, 54, and 50 acyl carbons were observed, each with a number of acyl double bonds. The predominate species was 52:2 (total acyl carbons:total acyl double bonds) representing ∼14 mol% of the TG class; and the combination of TG(52:3), TG(56:8), TG(54:3), TG(56:7), and TG(50:1) species represented 39% of TGs. The DG class was represented by 30 molecular species (29 detected in all three samples) (**Figure 6C**). Three principal groups with 32, 34 and 36 acyl carbons were identified, none of which contained double bonds. Approximately 58% of the DG pool was composed of the DG(32:0), DG(34:0), and DG(36:0) species. The cholesteryl ester class was composed of 20 molecular species (all species were detected in all samples) (**Figure 6D**) with four principal species: CE(18:1), CE(20:4), CE(22:4) and CE(22:5) making up 60% of cholesteryl esters.

The luteal LD phospholipid composition (expressed as a percentage of total phospholipid) was predominately PC (37%) followed by SM (26%), PI (15%), PE (13%), PS (7%) and PG (0.8%) while lysophospholipids only accounted for 1.4%. The phospholipid composition of luteal LD was surprisingly complex with more than 120 molecular species (85 detected in all three samples) in 5 classes (**Figure 6E**). Analysis detected a total of 31 PC molecular species (22 detected in all three samples), with PC(34:1) contributing to 43% of the population and an additional 33% of the PC class was made up of PC(36:4), PC(38:4), and PC(38:5). Lipidomics detected 34 species of PE (27 detected in all samples) which were dominated by PE(38:4), PE(36:2), PE(36:4), and PE(34:1). Twenty species of PI were detected (16 in all three samples) with 72% of the total PI pool represented by PI(38:4), PI(40:5), and PI(38:5). Lipidomics detected 18 species of PS (13 in all three samples) with 76% of the total PS pool represented by PS(40:5), PS(40:4), PS(36:1), and PS(38:3). The PG class was composed of 14 species (7 detected in all samples) with PG(36.2) contributing to 46% of the total PG pool.

## 6. Discussion

Luteal LDs are a prominent feature of bovine luteal tissue and the presence, prominence, and composition of luteal LDs distinguish luteal tissue and luteal cells from other tissues and cell types. The large and small steroidogenic luteal cells contain these distinguishing LDs; with large luteal cells containing numerous small LDs, and small luteal cells processing fewer, but larger, LDs. Luteal LDs have a diameter of 200 nm – 1100 nm, confirmed by a variety of techniques including transmission electron microscopy, confocal microscopy, and phospholipid: total lipid ratio. Purified luteal LDs are rich in in TG and contain cholesteryl esters, supporting the concept that luteal LDs have an important role in cellular metabolism and steroid biosynthesis by storing steroidogenic precursors.

Luteal tissue of all examined species contain abundant LDs which vary in size and number throughout the normal estrous or menstrual cycle (15–19). The findings that neutral lipids occupy approximately 5-16% of bovine luteal tissue area is in agreement with other studies examining LD to luteal cell area or volume (27, 28, 43). In contrast, other tissues rarely contain the same amount of neutral lipids, except during pathological situations (8–11). Likely, the area occupied by LDs in steroidogenic cells is even greater since luteal endothelial cells account for nearly 50% of luteal tissue (44) but rarely contain LDs (45). The present study and previous ultrastructural studies have noted the close contact of LDs with mitochondria (17, 21, 23). The proximity of luteal LDs to mitochondria suggests that these organelles may function cooperatively to synthesize steroids (6, 13). The dynamics of LD and mitochondria interactions in luteal cells are unknown, but recent studies in other tissues implicate the mitochondrial targeting regions of PLIN5 (46) and DGAT2 (diacylglycerol O-acyltransferase 2) facilitate physical interactions between mitochondria and LDs (47). The emerging field of mitochondrial associated membranes offers the possibility that there are regionalized areas of contact between mitochondria, endoplasmic reticulum and LDs (48, 49).

The size, distribution, protein, and lipid content of LDs can differ among various tissues (7, 13); therefore, expression of LD-associated proteins in bovine luteal tissue was compared with cardiac, hepatic, and adipose tissues. The tissues differed in the relative composition of PLINs and other lipid modifying enzymes, consistent with previous reports for adipose (50), cardiac (51), and hepatic tissue (10). Luteal tissue expressed PLIN2, PLIN3, PLIN5, hormone sensitive lipase, and ABHD5; and lower levels of ATGL and SOAT proteins; whereas, PLIN1 was undetectable. The presence of PLIN2, PLIN3, and PLIN5 suggest luteal LDs are metabolically active, oxidative, and mitochondrial-associated (7). As well, bovine tissues differed in lipid composition from other bovine tissues. Bovine luteal tissue contained higher levels of TG than other tissues (cardiac, hepatic, & pulmonary) with the unsurprising exception of adipose tissue. The high TG content of luteal tissue has unknown importance but the re-esterification of fatty acids liberated from cholesteryl esters during steroidogenesis could prevent fatty acid induced lipotoxicity. The unique LD-associated protein and lipid content of luteal LDs suggests luteal LDs have specialized lipid use and storage needs compared to other tissues.

After ovulation, the granulosa and theca cells of the follicle differentiate into the large and small steroidogenic cells, respectively, of the corpus luteum (34, 41). In contrast to the preovulatory follicle, which contains few LDs (24, 52), luteal tissue possesses numerous LDs. In this study, luteal tissue had fully established luteal LD content by three days post-ovulation, consistent with the idea that ovulation induces the formation of LDs during differentiation of the ovarian follicle into luteal tissue. Additionally, both large luteal cells and small luteal cells have abundant LDs in comparison to either granulosa or theca cells. These observations are in keeping with studies by Guraya *et al.* who described that following the luteinizing hormone surge, human granulosa cells develop fine “lipid granules” and “heterogeneous lipid bodies” within newly ruptured follicles, and that the theca interna cells of newly ruptured follicles fill with sudanophilic lipids, including cholesterol and cholesterol esters (24). As well, LD numbers and size increase in granulosa cells within a day after ovulation in rabbits (52). This study revealed that when compared to follicular cell types, bovine luteal cells have more protein and mRNA content of LD-associated proteins, PLIN2 and hormone sensitive lipase, paralleling the increase in LD formation. In keeping with the increase in luteal LDs, large and small luteal cells contain more TG than granulosa or theca cells. Similarly, treatment of rhesus macaques with luteinizing hormone increases the amount of PLIN2 protein in granulosa cells within 12 hours (22). The program of cellular differentiation following ovulation resulting in luteinization of granulosa and theca cells is accompanied by increased expression of LD-associated proteins, elevation in TG, and formation of LDs in steroidogenic luteal cells.

The LDs of large and small luteal cells differ primarily in the number and size of LDs per cell, with minimal differences in lipid content and LD-associated mRNAs. Small luteal cells have a mean LD volume 2.4 times greater than large luteal cells; whereas, large luteal cells have three times the number of LDs present in small luteal cells. This finding is in agreement with data by Khanthusaeng et al, who found that ovine large luteal cells have many smaller LDs and small luteal cells have fewer, but larger LDs (43). The difference in LD size and number in the two cell types may relate to the different functions of large and small luteal cells. Bovine large luteal cells are responsible for secretion of large quantities of progesterone under basal conditions; whereas, small luteal cells secrete less progesterone under basal conditions but respond acutely to luteinizing hormone with robust increases in progesterone production (53). Small LDs in large luteal cells could provide increased accessibility for continuous substrate utilization, such as liberation of cholesterol for constitutive steroid synthesis (7). The larger LDs seen in small luteal cells could function primarily as a lipid storage mechanism, which can be accessed as needed, such as in response to hormones that stimulate steroidogenesis (7).

Both TGs and cholesteryl esters are almost exclusively found in tissue LDs. Bovine luteal LDs contain primarily TG (89% of total lipid content), and are relatively cholesteryl ester-poor (3.7 ± 0.7 pmol/μg protein; 2% of total lipid), finding similar to whole bovine luteal tissue (25). Although studies (54) have shown that cholesteryl esters and TG can segregate into distinct LD populations; the LDs in either large or small luteal cells show little segregation of TG and cholesteryl esters, which may reflect species or cell type differences. In contrast to bovine, luteal tissue from rabbits and rats is cholesteryl ester-rich (26). Because bovine luteal tissue does not rely on de novo cholesterol synthesis for progesterone synthesis (55), a small pool of cholesteryl esters may be sufficient for luteal function. The high TG content in bovine luteal LDs may serve as a substrate for energy production to fuel the steroidogenic output of luteal tissue. The fatty acids derived from cholesteryl esters may be converted to biologically active lipid mediators, re-esterified and stored in LDs or cell membranes, or used for β-oxidation, ultimately producing acetyl-CoA for the citric acid cycle. It seems likely that the production of large quantities of progesterone by luteal cells could require β-oxidation of fatty acids to provide the energy needed for optimal steroidogenesis.

Mass spectrometry was used to obtain detailed information about the lipid composition of bovine luteal tissue LDs. LDs are composed of an core comprising of neutral lipids (TGs and cholesteryl esters) bound by a monolayer of phospholipids (7). Recent studies have exposed the dynamic lipid composition of LDs within numerous cell types and determined that the phospholipid composition of isolated droplets is remarkably complex. Luteal LD phospholipid composition (expressed as a percentage of total phospholipid) of luteal LDs is predominately PC (45%) followed by SM (22%), PI (13%) and PE (11%). In contrast phospholipid content of hepatic LD from fed animals was largely PC (61%) and PE (23%) with lesser amount of PI and SM (56). The present results also differ somewhat from a report on the phospholipid content of LDs isolated from various cells lines following incubation with oleate to induce LD formation; the LD were predominately PC rich followed by PE and PI, as noted above, but were deficient in SM and PS (56–58). When directly compared, luteal LDs had ∼15-fold more SM and ∼8.5-fold more PS than either mouse hepatic tissue or CHO K2 cells (56–58). Despite the tissue and cell type differences in lipid class abundance analysis of the specific TG, DG, PC, PE and PI species revealed that the constituent fatty acids within each phospholipid class are similar to previous studies (56–58).

In steroidogenic tissues, the hydrolysis of cholesterol esters by hormone sensitive lipase yields cholesterol and a free fatty acid. The cholesterol serves as a precursor for mitochondrial steroid synthesis (59), and while the fate of the fatty acid is unknown it could be converted to biologically active lipid mediators, re-esterified and stored in LDs or membranes, or used for energy production. The cholesteryl esters in bovine luteal LDs are primarily (60%) mono- and polyunsaturated fatty acids evenly distributed among oleic acid, and the omega-3 and omega-6 20:4, 22:4 and 22:5 fatty acids. This differs from ovine luteal cholesterol ester fatty acids which are predominantly palmitic acid [(16:0), 30.7%], oleic acid [(18:1) 22.3%] and linoleic acid [(18:2) 17.5%](16). Cholesteryl ester fatty acid content of rat luteal tissue had many similarities to those in bovine luteal LDs but contained more palmitic acid (16:0), and less oleic and arachidonic acids (18:1 and 20:4) content, and a reversed ratio of 18:0 to 18:1 fatty acids. The differences in fatty acid composition among these species could be a reflection of species or diet effects (15).

This study described in detail the extent, size, number and content of LDs in bovine luteal tissue and steroidogenic cells. The study also examined the increases in LD-associated proteins and lipids that occur during the follicle to luteal transition. Further research examining the impact of obesity, undernutrition and polycystic ovary syndrome on luteal LDs may provide insights into mechanisms of infertility and disorders of steroidogenesis. Additional studies examining the LDs in theca and granulosa cells and the mechanisms controlling the onset of LD presence in luteal cells could reveal their role in steroid production. Luteal LDs likely play a critical role in progesterone production by storing cholesteryl esters and interacting with mitochondria and endoplasmic reticulum to optimize steroid synthesis and provide substrates for energy production.

